# Developing all-in-one virus-like particles for Cas9 mRNA/single guide RNA co-delivery and aptamer-containing lentiviral vectors for improved gene expression

**DOI:** 10.1101/2022.01.19.476334

**Authors:** Manish Yadav, Anthony Atala, Baisong Lu

**Affiliations:** Wake Forest Institute for Regenerative Medicine, Wake Forest University Health Sciences, Winston-Salem, NC, U.S.A.

**Keywords:** Lentiviral vector, Virus-like particles, CRISPR/Cas9, Aptamer, ABP

## Abstract

Lentiviral vectors (LVs) are widely used for delivering foreign genes for long-term expression. Normal LVs contain the RNA genome, which is reverse transcribed into DNA and integrates into the host genome to mediate long-term gene expression. Recently, virus-like particles (VLPs) were developed for mRNA delivery. In these VLPs, packaging mRNA into the particles is achieved via interactions between the aptamer and aptamer-binding protein (ABP), and mRNA is not reverse transcribed or integrated. These VLPs are useful for delivering Cas9 mRNA for short-term endonuclease expression. Generating high-titer normal integrating LVs is challenging. Until recently, VLPs were not efficient for co-delivering Cas9 mRNA and single guide RNA (sgRNA). By fusing ABP to the N-terminus of HIV Gag protein, hybrid particles were developed to co-deliver Cas9 mRNA and sgRNA. But the method for modifying Gag impaired particle assembly. Previously we found that adding an ABP after the second zinc finger domain of nucleocapsid (NC) protein had minimal effects on particle assembly. Based on this observation, we developed hybrid particles to co-deliver Cas9 mRNA and sgRNA. We further improved LVs for integrated gene expression by including an aptamer sequence in lentiviral transfer plasmids, which improved lentiviral particle production and enhanced LV genomic RNA packaging. In summary, we describe development of new all-in-one VLPs for co-delivery of Cas9 mRNA and sgRNA and new LVs for enhanced vector production and gene expression.

## 1. Introduction

Lentiviral vectors (LVs) are widely used as gene delivery vectors in basic research and clinical applications. The advantages of LVs for gene transfer include long-term target gene expression, ability to transduce non-dividing cells, and relative low host immune responses to the vectors. One of the limitations of LVs is their relative low titer, which poses challenges to LV production. Efforts were made to improve LV production by increasing packaging cell transfection efficiency [1], improving transcription of genes coding for packaging viral proteins [2], blocking innate antiviral responses in packaging cells [3], and modifying the packaging cell culture conditions [4, 5]. These efforts improved LV production somewhat, but LV production is still a challenge for clinical applications.

The target expression levels of LVs are positively related to viral genome copy numbers per cell in the range of 1 to 4 [6, 7]. To balance gene expression and safety, the FDA recommends that the integration copy number be <5 copies per genome [8]. Each LV particle contains two copies of LV genomic RNA due to the enhanced affinity of the HIV-1 NC protein with the dimeric viral genome RNA [9]. Thus, increasing viral genomic RNA copy numbers per particle may increase target gene expression without causing safety concerns. However, so far, attempts to increase viral genomic copy numbers per particle via increasing dimerization of genomic RNA have not been successful [10].

CRISPR/Cas9 (clustered regularly interspaced short palindromic repeats/CRISPR-associated 9) endonucleases [11] create double-strand DNA breaks at specific genome loci [12-15], but also cleave off-target areas [16, 17]. Delivering Cas9 ribonucleoprotein (Cas9 RNP) [18] or Cas9 mRNA [19] enabled short-term expression of CRISPR/Cas9 and greatly improved safety. Based on these observations, viral vectors have been modified for delivering Cas9 RNPs [20-27] or Cas9 mRNAs [28-34] to improve safety.

Compared with Cas9 RNP delivering VLPs, Cas9 mRNA-delivering VLPs might achieve higher Cas9 protein expression and thus higher genome editing activity when targeting loci that are hard to access [35]. However, current Cas9 mRNA delivering VLPs either could not co-deliver Cas9 mRNA and sgRNA [28-30, 32], had inconsistent Cas9 mRNA and sgRNA co-delivery efficiency [31], or unknown particle assembly efficiency [33, 34].

We took advantage of interactions between RNA aptamers and ABPs to package and deliver Cas9 mRNA or RNPs by VLPs [24-26, 28, 29]. Contrary to fusing ABP to the N-terminus of Gag [33, 34] or removing NC [31], which affected particle assembly [20, 36], we inserted ABPs after the second zinc finger domain of the NC and observed minimal effects on particle assembly [24, 25, 28]. Here we report the development of all-in-one VLPs for co-delivery of Cas9 mRNA and sgRNA, and modified LVs to improve LV genomic RNA packaging and gene delivery.

## 2. Materials and Methods

### 2.1. Plasmids

pRSV-Rev (Addgene #12253), a plasmid conferring expression of HIV-1-derived Rev (coding for sequence-specific RNA-binding protein) under the transcriptional control of the Rous Sarcoma Virus (RSV) promoter; pMD2.G (Addgene #12259), a plasmid expressing the spike G glycoprotein of the vesicular stomatitis virus (VSV-G) for pseudotyping; pMDLg/pRRE (Addgene #12251), a third generation packaging plasmid; psPAX2-D64V (Addgene #63586), a second generation packaging plasmid with an inactivating D64V substitution in the integrase to prevent integration of the LV genomic RNA [37]; pLH-sgRNA1 (Addgene #75388), a lentiviral transfer plasmid for expressing *Streptococcus pyogenes* Cas9 (SpCas9) sgRNA; were purchased from Addgene and have been described previously. pLVX-IRES-ZsGreen1, a lentiviral transfer plasmid expressing *Zoanthus sp*. green fluorescent protein (ZsGreen), was purchased from Takara Bio Inc. (Kusatsu, Shiga, Japan). CmiR0001-MR03, a lentiviral transfer plasmid expressing green fluorescent protein (GFP), was purchased from GeneCopoeia, Inc. (Rockville, MD). Mammalian expression plasmid vector pcDNA3 was from Thermo Fisher Scientific (Waltham, MA). pFCK-HBB(n)-g1, a plasmid for expressing *Staphylococcus aureus* Cas9 (SaCas9) sgRNA targeting the mutant sequence causing sickle cell disease [38], and pSaCas9-HBB-sgRNA1^Tetra-com^, a plasmid expressing Cas9 and com-containing sgRNA targeting the *HBB* sickle mutant sequence [24], have been described previously. Other plasmids were generated for this study as described in **S1 Table**. Gene synthesis was done by GenScript Inc. All constructs generated were confirmed by Sanger sequencing. Sequence information for primers and oligonucleotides are listed in **S2 Table**. SpCas9 and SaCas9 target sequences and the oligonucleotides used for making the sgRNA expression constructs are listed in **S3 Table**.

### 2.2. Cell culture

HEK293T (ATCC CRL-11268™), Neuro-2A (ATCC CCL-131™) cells and GFP-reporter cells (see the following section for details) described previously [39] were cultured in Dulbecco’s Modified Eagle Medium (DMEM) with 10% fetal bovine serum (FBS), 100 U/ml penicillin, and 100 μg/ml streptomycin (Thermo Fisher Scientific) at 37°C in an incubator with 5% CO_2_.

### 2.3. Detecting gene editing activities by GFP-reporter assays

The HEK293T-derived GFP-reporter cells were used for detection of CRISPR/Cas9-induced insertions and deletions in the hemoglobin subunit beta (*HBB*) sickle mutant sequence and interleukin 2 receptor subunit gamma (*IL2RG*) target sequences [39]. The cells contained a GFP-reporter cassette, which has a 119 base-pair (bp) insertion right after the start codon of GFP cDNA to disrupt the GFP reading frame. This inserted sequence contained target sequences for the *HBB* sickle cell mutation and *IL2RG*. Without gene editing, GFP was not expressed because of the disrupted reading frame. With gene editing in either the *HBB* sickle mutation or the *IL2RG* target sequence, deletions of 3N+2 or insertions of 3N+1 base-pairs can restore GFP expression. GFP-positive cells were analyzed by a fluorescence microscope or an Accuri C6 flow cytometer (BD Biosciences, San Jose, CA) as described previously [28]. Genome editing activity was also assayed by targeted next generation sequencing (NGS), as described in the following sections.

### 2.4. Production of LVs and Cas9 mRNA- or Cas9 RNP-containing VLPs

lentiviral transfer plasmid pLH-sgRNA1-IL2RG, pFCK-HBB(n)-g1, pLVX-IRES-ZsGreen1 and pCSII-hCLCN5 were used to produce LVs for expressing *IL2RG-*targeting sgRNA, *HBB* sickle mutant sequence-targeting sgRNA, ZsGreen, and chloride voltage-gated channel 5 (CLCN5), respectively. The second and third generation packaging systems were used to package LVs as described previously [28]. Packaging plasmid psPAX2-D64V was used for packaging integration-defective LV (IDLV). Cas9 mRNA or Cas9 RNP VLPs were prepared as described previously [24, 28]. Co-packaging of Cas9 mRNA and sgRNA-expressing LV genomic RNA into VLPs were achieved by co-transfection of the following five plasmids (DNA amounts are listed for transfecting 6×10^5^ HEK293T cells cultured in 6-well plates): psPAX2-D64V (2 μg); psPAX2-D64V-NC-ABP (1 μg, see **Table 1** for specific psPAX2-D64V-NC-ABP packaging plasmids used); aptamer-modified Cas9 mRNA expression plasmid (2.25 μg, see **Table 1** for corresponding aptamer-modified Cas9 mRNA expression plasmids used); sgRNA-expressing lentiviral transfer plasmid (2.25 μg); and pMD2.G (1.5 μg). The four different ABPs were <u>M</u>S2 <u>c</u>oat <u>p</u>rotein (MCP, binding to RNA aptamer MS2) [40], <u>P</u>P7 <u>c</u>oat <u>p</u>rotein (PCP, binding to RNA aptamer PP7) [41], λ N22 peptide (binding to RNA aptamer BoxB) [42], and <u>C</u>ontrol <u>o</u>f <u>m</u>om (Com, binding to RNA aptamer com) [43, 44]. MCP, PCP, N22 and Com expressed from the packaging plasmids will bind to the corresponding MS2, PP7, BoxB and com aptamers expressed from the aptamer-modified Cas9 mRNA expression plasmids, which explains the packaging of Cas9 mRNA into the VLPs.

**Table 1.** ABP-modified packaging plasmids and corresponding aptamer-modified Cas9 mRNA expression plasmids used for packaging Cas9 mRNA.

| Aptamer/<br>ABP pair | ABP-modified<br>packaging<br>plasmid | Aptamer-modified Cas9 mRNA expression plasmid |  |
| --- | --- | --- | --- |
|  |  | For SaCas9 mRNA | For SpCas9 mRNA |
| MS2/MCP | psPAX2-D64V-NC-MCP | pSaCas9<sup>1xMS2</sup>-2x3'UTR | pSpCas9<sup>1xMS2</sup>-2x3'UTR |
| PP7/PCP | psPAX2-D64V-NC-PCP | pBlue-SaCas9-3 aptamers | pBlue-SpCas9-3 aptamers |
| BoxB/N22 | psPAX2-D64V-NC-N22 | pBlue-SaCas9-3 aptamers | pBlue-SpCas9-3 aptamers |
| com/Com | psPAX2-D64V-NC-Com | pSaCas9<sup>1xcom</sup>-2x3'UTR or<br>pBlue-SaCas9-3 aptamers | pspCas9<sup>1xcom</sup>-2x3'UTR<br>pBlue-SpCas9-3 aptamers |

FuGENE HD (Promega, Madison, WI) or polyethylenimine (PEI, SynChem Inc., Elk Grove Village, IL) was used for DNA transfection at a ratio of 3 μl FuGENE HD or 2 μl PEI (1 mg/ml) for 1 μg DNA. Specifically, 6×10^5^ HEK293T cells were seeded in 6-well plates the day before transfection. Right before transfection, the cells were changed into 1 ml Opti-MEM. The DNA mixture were added to 0.2 ml Opti-MEM, followed by 27 μl FuGENE HD or 18 μl PEI. The DNA and FuGENE HD or PEI mixture were incubated at room temperature for 15 min, and were then added to the cells in Opti-MEM. The medium was changed to 2 ml Opti-MEM 24 h after transfection, and the LVs or VLPs were collected 72 h after transfection. The supernatant was spun for 10 min at 500 g to remove cell debris. The cleared supernatant was used for transducing cells.

### 2.5. LV and Cas9 VLP quantification

Concentrations of LVs and Cas9 VLPs were determined by LV-associated HIV p24 capsid protein (CA)-based ELISA (QuickTiter™ Lentivirus Titer Kit, Catalog Number VPK-107, Cell Biolabs, Inc. San Diego, CA). LVs or VLPs were precipitated according to the manufacturer’s instructions so that soluble p24 protein was not detected.

### 2.6. RNA isolation and RT-qPCR analyses

A miRNeasy Mini Kit (Cat No. 217004, QIAGEN, Germantown, MD) was used to isolate RNA from 150 μl unconcentrated particles (∼15 ng p24) or ∼10^5^ transduced cells. The QuantiTect Reverse Transcription Kit (QIAGEN) was used to reverse-transcribe the RNA to cDNA. Primers specific for codon-optimized *CLCN5* (hCLCN5-CF and hCLCN5-CR) were used to detect human *CLCN5* gene expression from the transgene. A custom designed sgRNA Taqman probe (Thermo Fisher Scientific, assay ID: APPRKYP; target sequence: gttttagtactctggaaacagaatctactaaaacaaggcaaaatgccgtgtttatctcgtcaacttgttggcgaga) was used for detecting *HBB* sickle mutant sequence targeting sgRNA. The *GAPDH* (Glyceraldehyde 3-phosphate dehydrogenase) Taqman probe (Thermo Fisher Scientific) was used as an endogenous control.

RT-qPCR was used to detect LV genomic RNA in particles with primers Psi-F and Psi-R. LVs with and without com in LV genomic RNA were assayed side by side. We set the genome copy number of LVs without com at two. Genome copy numbers of LVs with com were calculated after normalizing particle input. Primer sequences are listed in **S2 Table**. SYBR Green and Taqman master mixtures (Thermo Fisher Scientific) were used for qPCR using a QuantStudio™ 3 instrument.

### 2.7. LV and VLP transduction

Unconcentrated LVs or VLPs (10∼40 ng p24 protein) were added to 2.5×10^4^ HEK293T or Neuro-2A cells grown in 24-well plates with 8 μg/ml polybrene. The cells were incubated with the particle-containing medium for 12-24 h, after which the medium was replaced with DMEM medium containing 10% FBS and antibiotics.

### 2.8. Next-generation sequencing and data analysis

Report-F and Report-R2 primers were used to amplify the target DNA sequence from GFP-reporter cells for targeted NGS (Amplicon-EZ, GENEWIZ, Morrisville, NC). The proofreading HotStart® ReadyMix from KAPA Biosystems (Wilmington, MA) was used for PCR. About 50,000 reads/amplicon were obtained. Insertions and deletions were analyzed with the online software Cas-Analyzer [45] and CRISPRESSO2 [46].

### 2.9. Statistical analysis

GraphPad Prism software (version 5.0) was used for statistical analyses. T-tests were used to compare the averages of two groups. Analysis of variance (ANOVA) was performed followed by Tukey multiple comparisons tests to analyze data from more than two groups. Bonferroni *post hoc* tests were performed following ANOVA in cases of two factors. P<0.05 was regarded as statistically significant.

## 3. Results

### 3.1. Fusing ABP to NC protein of Gag impaired LV gene transfer

Several studies have found that naked sgRNAs could not be functionally delivered by VLPs [24, 31, 32], possibly because the unprotected sgRNAs were unstable when escaping from the endosome system in the recipient cells. We wondered whether VLPs could be developed for co-delivering Cas9 mRNA and sgRNA. In recipient cells, the Cas9 mRNA molecules will serve as the translation templates for short-term Cas9 protein expression, yet will not be reverse transcribed or integrated. The sgRNAs will be expressed from the sgRNA-expressing LV genomic RNA co-packaged with Cas9 mRNA into the particles. The LV genomic RNA will be reverse transcribed into DNA for sgRNA expression, but will not be integrated due to the D64V mutation in the integrase of the particles. The idea was to package Cas9 mRNA via aptamer and ABP interactions, and to package the sgRNA-expressing LV genomic RNA by its lentiviral packaging signal (Psi). To package Cas9 mRNA via ABP interactions, we inserted ABPs after the second zinc finger domain of NC [24, 28]. Although this modification showed minimal effects on lentiviral particle assembly [24, 28], its effects on LV gene transfer were unclear. Thus, we first tested whether our way of modifying the packaging plasmid [24, 28] affected LV-mediated gene expression.

We generated GFP-expressing IDLVs with MCP- or Com-modified packaging plasmids, and transduced the resulting IDLVs into HEK293T cells. Inserting MCP or Com into this position of Gag reduced the percentage of GFP-positive cells more than 20 times compared with IDLVs generated with packaging plasmid without ABP modification (**Fig 1A**). This observation was confirmed by independent repeated experiments (**S1 Fig**). Since Com or MCP modification of the packaging plasmid had little effect on lentiviral particle assembly [24, 28], the marked decrease of GFP-positive cells could not be explained by different particle production efficiency. Thus, fusing ABP after the second zinc finger domain of NC greatly diminished gene expression from the modified LVs.

**Fig. 1.**
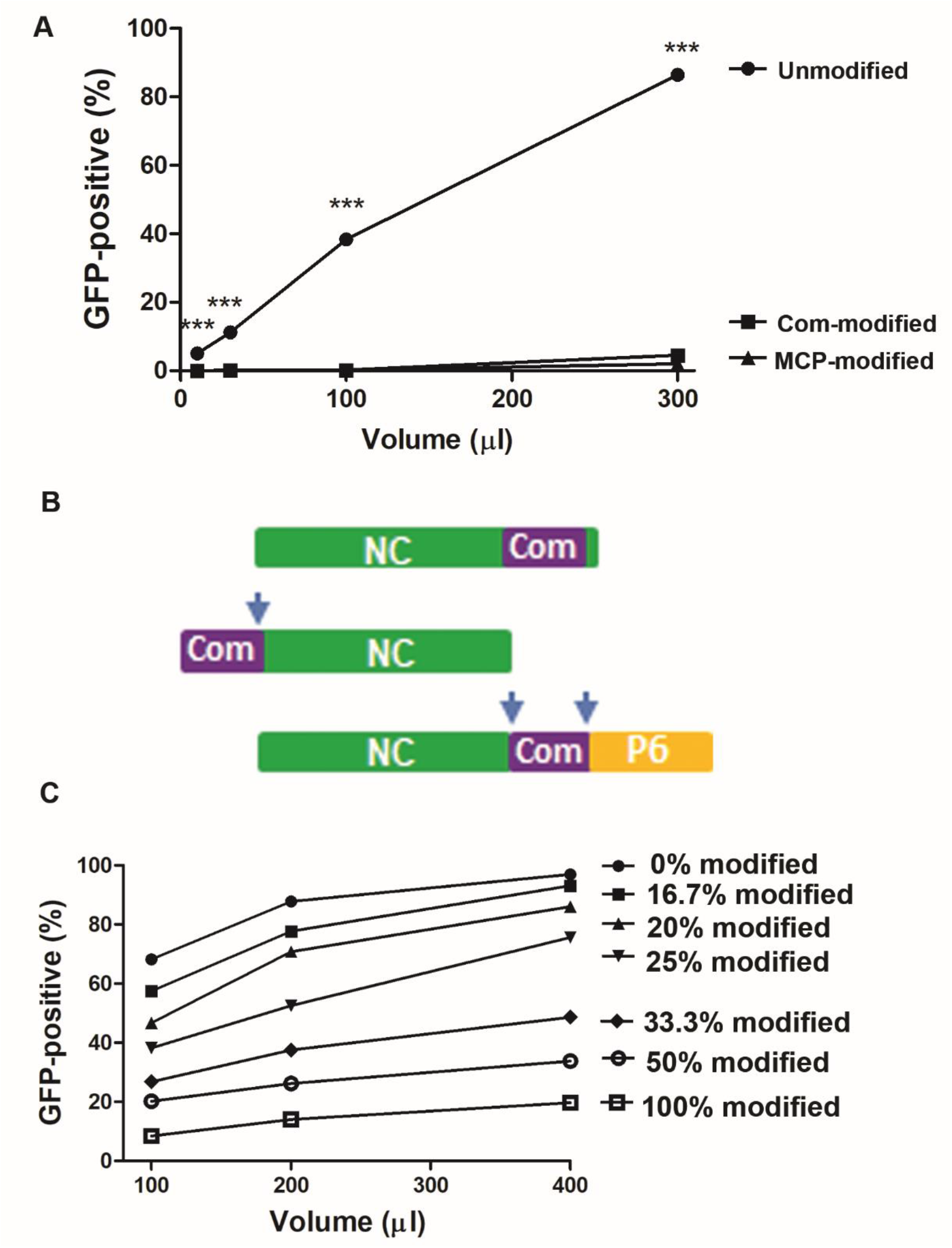
Rescuing LV titer by unmodified package plasmid DNA. **A**. Inserting Com or MCP after the second zinc finger domain greatly affected LV titer. Com- or MCP-modified packaging plasmid was used to generate GFP-expressing IDLVs. *** indicates p<0.0001 compared with other LVs of the same volume (n=3, Bonferroni *post hoc* tests following ANOVA). Unmodified, Com-modified, and MCP-modified package plasmids were psPAX2-D64V, psPAX2-D64V-NC-Com, and psPAX2-D64V-NC-MCP, respectively. **B**. Different Gag modifications tested. ABP (MCP or Com) was fused after the second zinc finger domain of NC (top row), fused at the N-terminus of NC (middle), or expressed between NC and P6. Arrows indicate protease cleavage sequences. **C**. Rescuing LV titer by unmodified packaging plasmid DNA. GFP-expressing IDLVs were made with different ratios of modified (psPAX2-D64V-NC-Com) and unmodified packaging plasmid DNA (psPAX2-D64V). For (**A** and **C**), indicated volumes of IDLV-containing supernatants were used to transduce 2.5×10^4^ HEK293T cells followed by flow cytometry analysis. Our typical LV-containing supernatants contained 100∼150 ng/ml p24. Means of 3 technical replicates are shown. Each experiment was performed at least twice with similar results (see Supplementary figures S1 and S2).

We also tested two more ways of Gag modification: 1) fusing ABP Com at the N-terminus of NC and 2) inserting Com at the C-terminus of NC with protease cleavage sites between Com and the adjacent viral proteins, so that separate viral proteins and Com could be generated by protease processing (**Fig 1B**). Efficient GFP expression from the resulting LVs was not achieved (unpublished data). Thus, fusing ABP with Gag to preserve both high particle assembly efficiency and gene transfer ability could not be achieved with the methods tested.

Since each lentiviral particle has 2500-5000 Gag precursors [47], we wondered whether LV gene transfer capability could be rescued by supplementing particles with unmodified Gag precursor protein. We thus prepared GFP-expressing IDLVs with different ratios of modified and unmodified Gag precursor using psPAX2-D64V-NC-Com and psPAX2-D64V, respectively. Increasing unmodified packaging plasmid DNA rescued GFP expression of the particles (**Fig 1C**). With only 20% modified packaging plasmid DNA, the resulting GFP-expressing IDLVs showed over 80% infection efficiency of IDLVs produced with 0% modified packaging plasmid DNA. Similar results were observed in an independent repeated experiment (**S2 Fig**). Since this method of modifying Gag precursors did not affect particle assembly, we concluded that supplementing particles with unmodified Gag precursor protein supported both particle assembly and vector genome transfer.

### 3.2. Finding the most efficient aptamer and ABP pair for Cas9 mRNA delivery

In our previous study [28], we used aptamer/ABP pairs MS2/MCP and PP7/PCP for Cas9 mRNA packaging. Currently, MS2/MCP is the most widely used aptamer/ABP pair for Cas9 mRNA delivery [28-34]. We recently compared four pairs of aptamer/ABP, including MS2/MCP, PP7/PCP, BoxB/λ N22 peptide, and com/Com, for VLP packaging of Cas9 RNPs, and com/Com was the most efficient pair for this purpose [24]. To determine whether com/Com was also more efficient than MS2/MCP for mRNA packaging, we compared these two pairs of aptamer/ABP for Cas9 mRNA delivery. We made a SaCas9-expressing construct (pSaCas9^1xcom^-2×3’UTR, see plasmid 11 in **S1 Table**) with one copy of a com aptamer replacing the MS2 aptamer of pSaCas9^1xMS2^-2×3’UTR that we published previously [28].

We packaged SaCas9 mRNA with packaging plasmid psPAX2-D64V-NC-MCP, expressing a Gag protein with MCP inserted after the second zinc finger domain of the NC protein for packaging MS2 containing SaCas9 mRNA, or packaging plasmid psPAX2-D64V-NC-Com, expressing a Gag protein with Com inserted after the second zinc finger domain of the NC protein for packaging com containing SaCas9 mRNA. We assayed the genome editing activities of the particles by transfecting plasmid DNA expressing the sickle sgRNA into the GFP-reporter cells we generated for assaying CRISPR/Cas9 genome editing activities targeting *HBB/IL2RG* target sequences (**S3 Fig**) [39].

The com/Com SaCas9 mRNA VLPs generated significantly more GFP-positive cells (reflecting genome editing activities) than the MS2/MCP Cas9 mRNA VLPs (**Fig 2B**). To find the most efficient aptamer/ABP pair for Cas9 mRNA delivery, we further compared com/Com, BoxB/N22 and PP7/PCP for this purpose. We made a SaCa9 mRNA expressing plasmid pBlue-SaCas9-3 aptamers (**S1 Table**) containing one copy of a PP7, BoxB and com aptamer after *HBB* 3’-UTR, and packaged SaCas9 mRNA with a Gag protein in which PCP, λ N22 peptide, or Com was inserted after the second zinc finger domain of the NC protein (See **Table 1** for plasmids used). The com/Com Cas9 mRNA VLPs generated more GFP-positive cells than the other VLPs (**Fig 2C**). Interestingly, the two com-containing SaCas9 expressing plasmids, pSaCas9^1xcom^-2×3’UTR (**Fig 2B**) and pBlue-SaCas9-3 aptamers (**Fig 2C**) had the same SaCas9 expression cassette. However, Cas9 mRNA VLPs prepared with pBlue-SaCas9-3 aptamers had much higher genome editing activities than those prepared with pSaCas9^1xcom^-2×3’UTR. Possible explanations include 1) pSaCas9^1xcom^-2×3’UTR but not pBlue-SaCas9-3 aptamers had adeno-associated virus inverted terminal repeats, which might affect DNA secondary structure and gene expression of transfected DNA; 2) pBlue-SaCas9-3 aptamers but not pSaCas9^1xcom^-2×3’UTR had PP7 and BoxB aptamers in addition to com aptamer and might make the com aptamer more stable or accessible. We thus used pBlue-SaCas9-3 aptamers for further study.

**Fig. 2.**
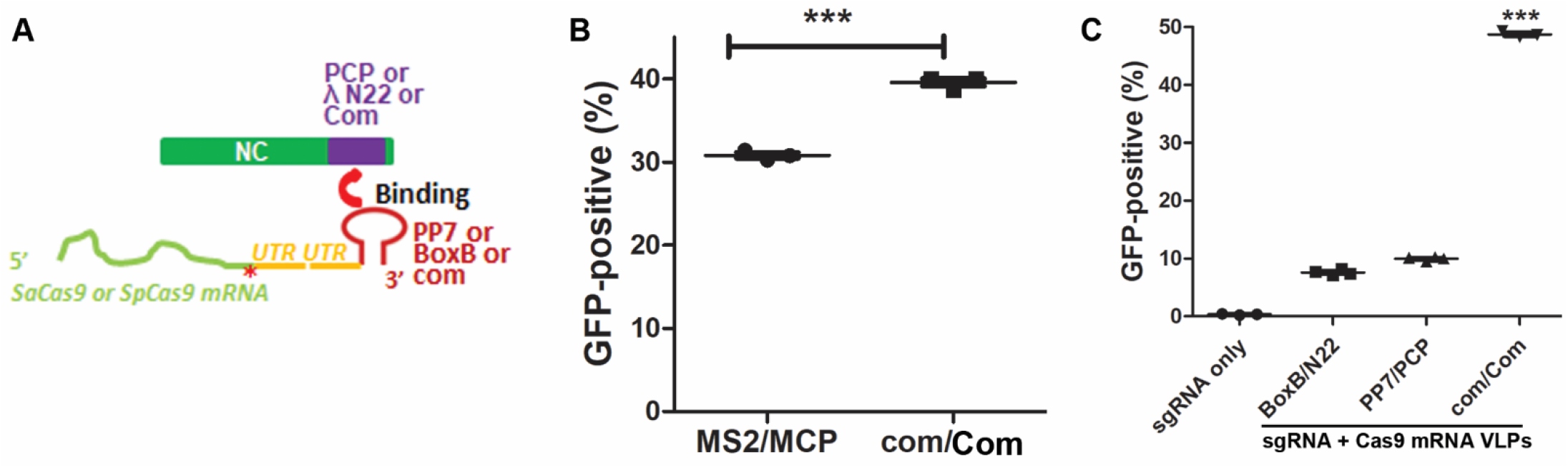
Finding the most efficient aptamer/ABP pair for Cas9 mRNA delivery. **A**. Aptamers and ABPs tested for Cas9 mRNA delivery by VLPs. PP7, BoxB, or com replaced MS2 in the Cas9-expressing construct and PCP, λ N22 peptide, or Com replaced MCP in the packaging plasmid. Red asterisk indicates stop codon. UTR: Human *HBB* 3’ untranslated region sequence for improving Cas9 mRNA stability and translatability. **B**. Comparing com/Com and MS2/MCP aptamer/ABP pairs for SaCas9 mRNA delivery. *** indicates p<0.0001 (two-tailed t-test). **C**. Comparing com/Com, BoxB/N22 and PP7/PCP aptamer/ABP pairs for SaCas9 mRNA delivery. *** indicates p<0.0001 for com/Com VLPs versus BoxB/N22 or PP7/PCP VLPs (Tukey’s *post hoc* analysis following ANOVA). For (B and C), 0.25 μg sickle sgRNA-expressing lentiviral transfer plasmid DNA was transfected into 2.5×10^4^ GFP-reporter cells 12 hours before transducing 500 μl indicated Cas9 mRNA VLPs. Our typical VLP-containing supernatants contained 100∼150 ng/ml p24

Similar to SaCas9 mRNA delivery, the com/Com pair was also most efficient for packaging and delivering SpCas9 mRNA (**S4 Fig**). Since our purpose was to find the most efficient aptamer/ABP pair for mRNA delivery, we did not adjust p24 to normalize particle input, but used the same volume of raw VLP-containing supernatant. The rationale is that genome editing activities of the VLPs are the combined results of multiple features: the efficiency of particle assembly after ABP insertion into NC, the capability of ABP to fold into a functional protein and to bind aptamer, the ability of the ABP to release the mRNA in recipient cells and the effects of the aptamer/ABP interaction on Cas9 mRNA stability and translatability. Thus the genome editing activity was the most relevant readout, and no single parameter (particle assembly, mRNA recruitment, mRNA release or mRNA stability and translatability) is able to predict the performance of an aptamer/ABP pair. Thus we believe that using equal particle volumes rather than particle numbers (p24) is the best way to compare different aptamer/ABP pairs. After finding that com/Com was more efficient than the most widely used MS2/MCP pair for Cas9 mRNA delivery, we used com/Com for subsequent all-in-one VLPs to co-deliver Cas9 mRNA and sgRNA.

### 3.3. VLPs can co-package Cas9 mRNA and sgRNA-expressing LV genomic RNA

We next asked whether Cas9 mRNA and sgRNA-expressing LV genomic RNA could be co-packaged into VLPs, with Cas9 mRNA packaged via aptamer (com) and ABP (Com) interactions and sgRNA-expressing LV genomic RNA packaged via Psi and NC interactions (**Fig 3A**). To test this idea, we co-packaged SaCas9 mRNA and LV genomic RNA, the latter contained an expression cassette to express a sgRNA targeting the mutant *HBB* sequence causing sickle cell disease (sickle sgRNA). VLPs were prepared by co-transfecting 5 plasmids into packaging HEK293T cells: Modified and unmodified packaging plasmids, VSV-G-expressing plasmid, Cas9-com-expressing plasmid, and sickle sgRNA-expressing lentiviral transfer plasmid.

**Fig. 3.**
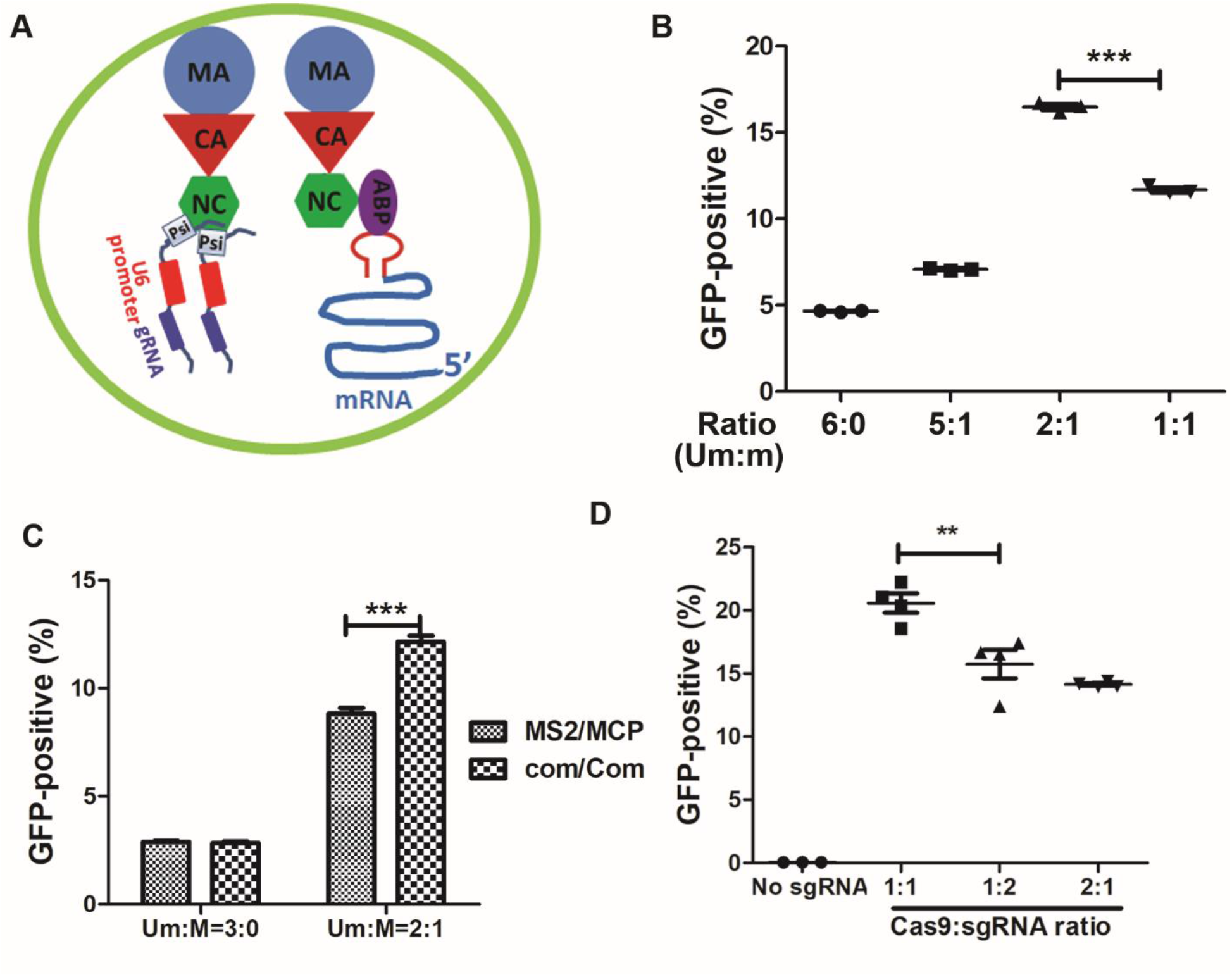
Co-packaging Cas9 mRNA and sgRNA-expressing LV genomic RNA in VLPs. **A**. Illustration of co-packaging mRNA and LV genomic RNA into VLPs. Envelope proteins are not shown. Only one modified and one unmodified Gag precursor in the immature virion are shown. **B**. Determining the best ratio of unmodified and modified packaging plasmid for most efficient co-packaging. Indicated ratios of unmodified (Um) packaging plasmid (psPAX2-D64V) and Com-modified (M) packaging plasmid (psPAX2-D64V-NC-Com) were used to transfect HEK293T cells by PEI to make VLPs, which contained Cas9 mRNA and LV genomic RNA for expressing sickle sgRNA. The particles (500 μl) were transduced into GFP-reporter cells and GFP-positive cells were analyzed by flow cytometry. Each data point indicates one technical replicate from the same batch of VLP particles. **C**. com/Com outperformed MS2/MCP for mRNA and sgRNA expressing cassette co-packaging. Experiments were done similarly to those in (**B**). **D**. Determining the best ratio of Cas9-expressing DNA and sgRNA-expressing lentiviral transfer plasmid DNA for most efficient co-packaging. Experiments were done similarly to those in (**B**), with a psPAX2-D64V and psPAX2-D64V-NC-Com ratio of 2:1. *** indicates p<0.0001, Tukey’s *post hoc* test following ANOVA (**B** and **D**) or Bonferroni *post hoc* tests following ANOVA (**C**). (**B-D**): 500 μl unconcentrated VLP-containing supernatant was added to 2.5×10^4^ GFP-reporter cells.

We titrated the unmodified and modified packaging plasmid DNA for co-packaging the Cas9 mRNA and sgRNA-expressing LV genomic RNA. Unconcentrated co-packaged VLPs were transduced into GFP-reporter cells to test for the presence of GFP-positive cells. The highest percentage of GFP-positive cells (reflecting genome editing activities) was observed when the unmodified versus modified packaging plasmid DNA were used at a 2:1 ratio (**Fig 3B**). Similar observations were made when VLPs were made by FuGENE HD-mediated transfection, which generated more active VLPs than when PEI was used for transfection (**S5 Fig**). Our results indicate that a Cas9 mRNA and sgRNA-expressing LV genomic RNA could be co-packaged in hybrid VLPs.

Consistent with packaging Cas9 mRNA only, com/Com also outperformed MS2/MCP for co-packaging Cas9 mRNA and sgRNA-expressing LV genomic RNA (**Fig 3C**). The observation of genome editing activities in VLPs where Cas9 mRNA was packaged via MS2/MCP interaction also suggested that the gene transfer function of MCP-modified packaging plasmid DNA could be rescued by unmodified packaging plasmid DNA. We further tuned the amount of Cas9 expression in DNA and sgRNA-expressing lentiviral transfer plasmid DNA, and found that the most efficient genome editing was achieved with equal amounts of each DNA (**Fig 3D**).

### 3.4. Cas9 mRNA and sgRNA-expressing LV genomic RNA co-existed in the same particle

Next, we tested whether the Cas9 mRNA and sgRNA lentiviral genome co-existed in the same particle. We prepared three types of particles in parallel: VLPs with Cas9 mRNA only, particles with *HBB* sickle sgRNA-expressing LV genomic RNA only (IDLV), and VLPs co-packaged with both. During transfection to generate the first two particles, equal amounts of non-relevant plasmid (pCDNA3) DNA were used to replace the sgRNA-expressing lentiviral transfer plasmid DNA for making Cas9 mRNA VLPs or the Cas9-expressing plasmid for making sgRNA-expressing IDLVs. Equal volumes of the two singly packaged particles (each 400 μl) were used to co-transduce GFP-reporter cells and to compare them with 400 μl co-packaged VLPs for genome editing. If the Cas9 mRNA and sgRNA-expressing LV genomic RNA did not co-exist in the same particle, then we would expect that a portion of the particles only contained Cas9 mRNA and the rest only contained the sgRNA-expressing LV genomic RNA. In this case, co-packaged VLPs should have lower genome editing activity than the combination of singly packaged particles. However, we found that co-packaged VLPs generated more GFP-positive cells (reflecting genome editing activities) than the combination of the two singly packaged particles (**Fig 4A**).

**Fig. 4.**
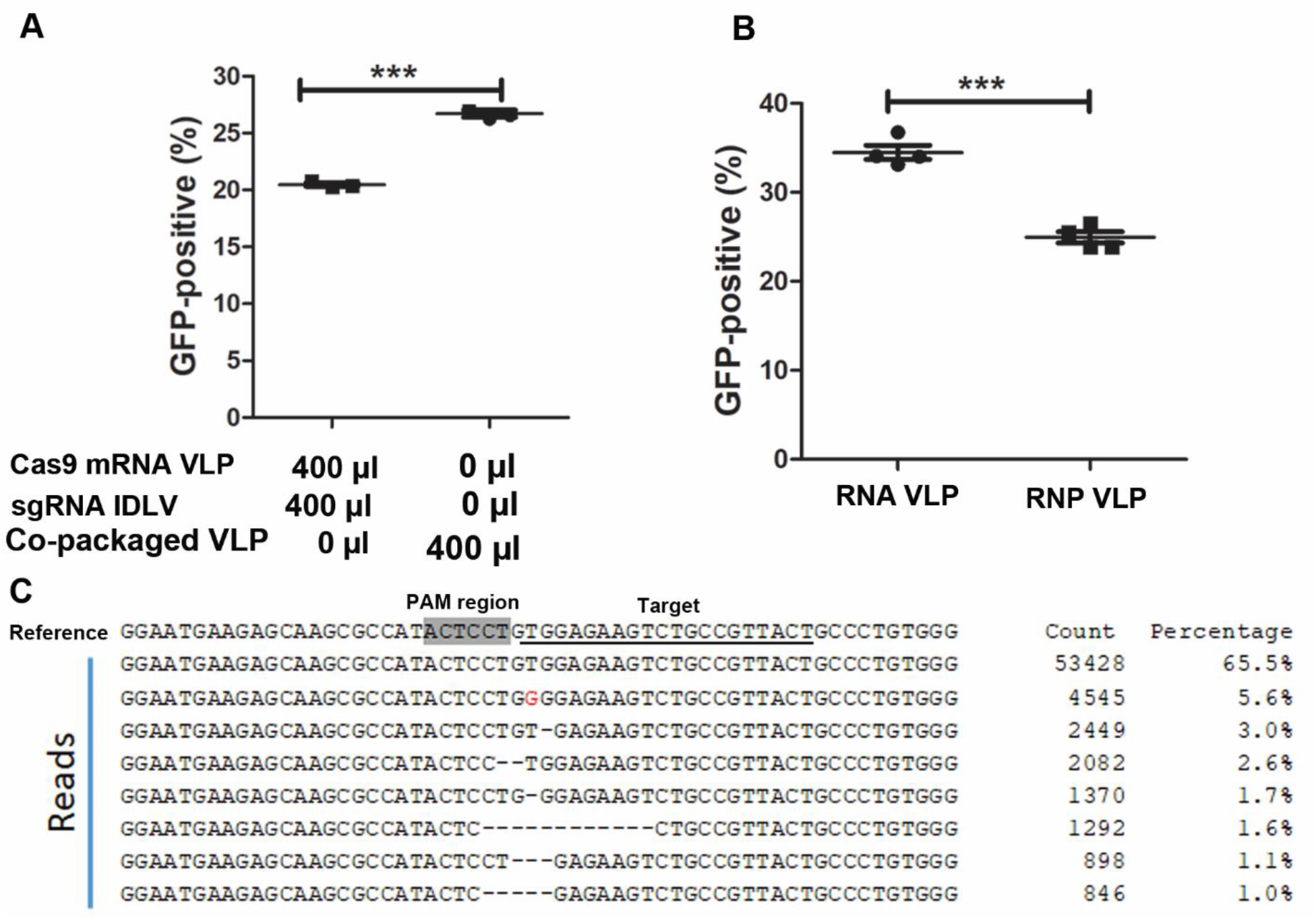
Characterization of Cas9 mRNA and sgRNA all-in-one VLPs. **A**. Genome editing activities of singly packaged particles (Cas9 mRNA VLPs or sgRNA IDLVs) versus co-packaged VLPs. For single packaging, either the Cas9-expressing DNA (pBlue-SaCas9-3 aptamers) or the sickle sgRNA-expressing lentiviral transfer plasmid DNA [pFCK-HBB (n)-g1] was replaced with pCDNA3 DNA to control transfection conditions. **B**. Genome editing activities of the co-packaged RNA VLPs and RNP VLPs. Five hundred microliter of supernatant were added to 2.5×10^4^ GFP-reporter cells. *** indicates p<0.0001, t-tests (**A, B**). Three preparations of VLPs were analyzed in each experiment. **C**. Deep sequencing analysis of insertions and deletions in the target region. The PAM (on the opposite strand) region is highlighted and the target sequence is underlined. The red “G” indicates substitutions. DNA was amplified from GFP-reporter cells treated with 50 μl all-in-one VLP-containing supernatant.

Ideally, observing the two types of RNAs in the same particle would provide definitive evidence of co-packaging. However, it is technically challenging to do so. Nevertheless, our results suggested that Cas9 mRNA and the sgRNA-expressing LV genomic RNA co-existed in the same particle. We thus call the VLPs with both Cas9 mRNA and LV genomic RNA all-in-one VLPs.

### 3.5. Cas9 mRNA and sgRNA all-in-one VLPs were more active than Cas9 RNP VLPs in genome editing

We further compared the genome editing activities of VLPs delivering co-packaged Cas9 mRNA and sgRNA-expression genomic RNA (RNA VLPs), and VLPs delivering Cas9 RNPs (RNP VLPs), both particles targeting the *HBB* sickle mutant sequence. RNP VLPs were prepared by co-transfection of three plasmids as we described recently [24]: the Com-modified packaging plasmid psPAX2-D64V-NC-Com, the Cas9 and com-containing sgRNA expressing plasmid pSaCas9-HBB-sgRNA1^Tetra-com^, and VSV-G expressing plasmid pMD2.G. We found that RNA VLPs (co-packaged with Cas9 mRNA and sgRNA-expressing LV genomic RNA) were more active in genome editing than RNP VLPs (**Fig 4B**). We analyzed the target DNA regions by targeted deep sequencing and observed 30% insertions and deletions in cells treated with the co-packaged RNA VLPs (**Fig 4C**). Note that this level of editing was achieved with only 50 μl un-concentrated particles (5∼10 ng p24) on 2.5×10^4^ cells, which is a very low dosage compared with similar studies [24, 28].

The Gag precursor protein expressed from psPAX2-D64V without ABP modification could also package Cas9-MS2 and Cas9-com mRNA, and produce detectable levels of genome editing (**Figs 3B** and **3C**), although the presence of ABPs greatly increased the genome editing activities. This observation suggested weak interaction between the lentiviral proteins and Cas9-aptamer mRNA.

### 3.6. Improving LV genomic RNA packaging with an RNA aptamer

The observation that mixing modified and unmodified Gag precursor supported relative efficient gene transfer activities of LVs prompted us to test whether the aptamer-ABP interaction could be used to increase LV genomic RNA packaging. We inserted the RNA aptamer com sequence between the Rev Response element (RRE) and central polypurine tract/central termination sequence (cPPT/CTS) of a ZsGreen-expressing lentiviral transfer plasmid (**Fig 5A**). This location was picked because it was outside of the sequences with known functions and the PpuMI restriction enzyme site was available for sub-cloning in most lentiviral transfer plasmids.

**Fig. 5.**
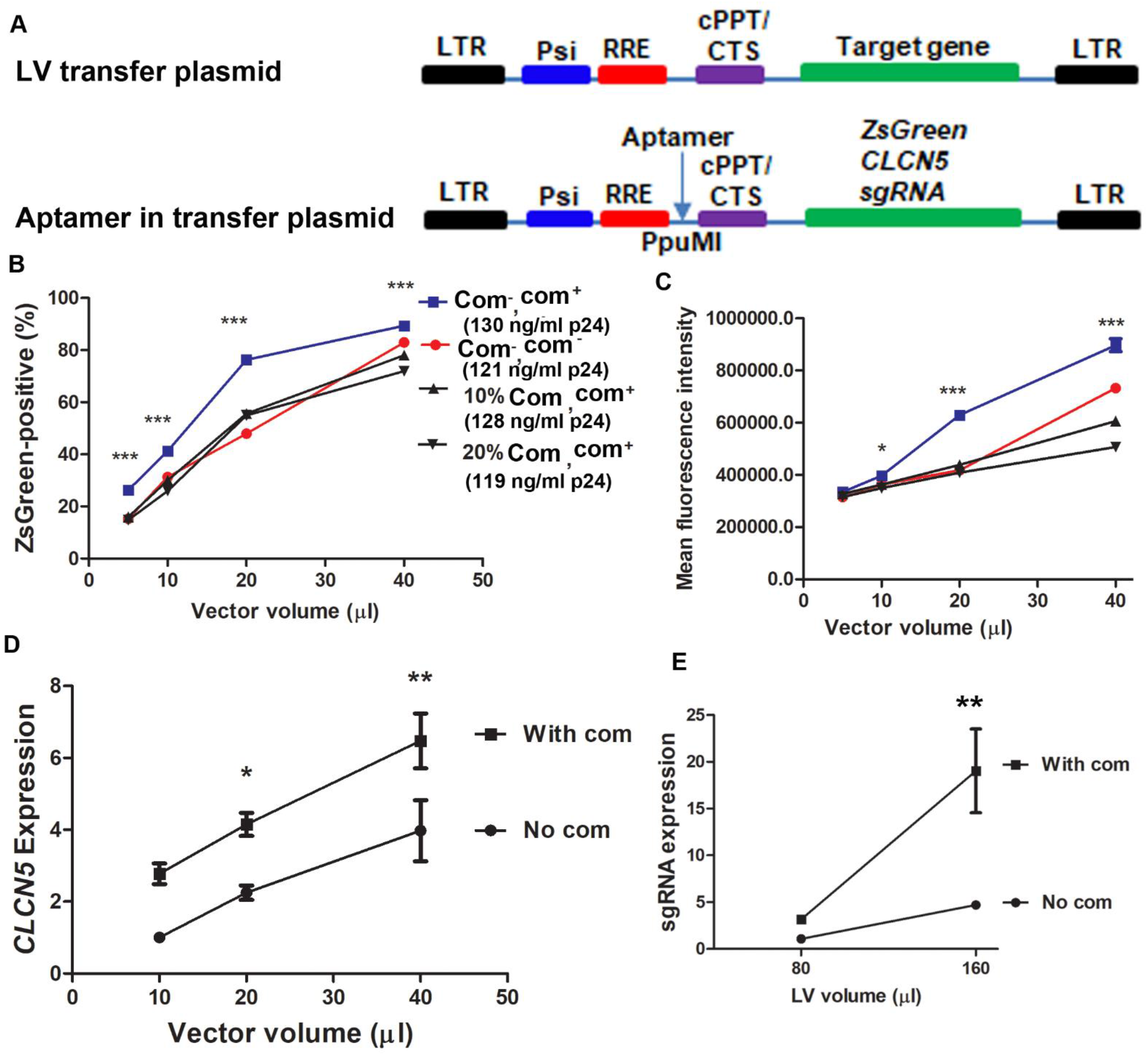
Including aptamer com in LV genomic RNA increases genomic RNA packaging and target gene expression. **A**. Illustration of common elements in lentiviral transfer plasmid (top) and the position of aptamer insertion (bottom). LTR: long terminal repeats; Psi: packaging signal; RRE: Rev Response element; cPPT/CTS: central polypurine tract/central termination sequence. **B**. Including com in LV genomic RNA increased the percentage of ZsGreen-positive cells. Com^-^,com^+^: packaging plasmid without Com modification and LV genomic RNA with com aptamer; Com^-^,com^-^: packaging plasmid without Com modification and LV genomic RNA without com aptamer; 10% Com,com^+^: packaged with 90% of unmodified and 10% Com-modified packaging plasmid DNA and LV genomic RNA with com aptamer; 20% Com,com^+^: packaged with 80% of unmodified and 20% Com-modified packaging plasmid DNA and LV genomic RNA with com aptamer. Representative LV concentrations determined by p24 ELISA are indicated in parentheses. ZsGreen-positive percentages were determined by flow cytometry. Both modified and unmodified packaging plasmid DNA had a D64V mutation in integrase. **C**. Including com in LV genomic RNA increased the intensity of ZsGreen signal. Symbols are the same as in (**B**). **D**. Including com in LV genomic RNA increased *CLCN5* gene expression. *CLCN5* expression was determined by RT-qPCR with primers specific to the codon-optimized *CLCN5* cDNA. **E**. Including com in LV genomic RNA increased *HBB* sickle sgRNA expression. 2.5×10^4^ HEK293T cells (**B, C** and **D**) or Neuro-2A cells (**E**) were transduced with indicated volume of each LVs. sgRNA expression was determined by RT-qPCR. *, ** and *** indicates p<0.05, p<0.01, and p<0.0001, in Bonferroni *post hoc* tests following ANOVA (**B, C, D** and **E**). Three biological replicates were included in all experiments. Typical particle concentration of the LVs was 100-200 ng p24/ml.

We generated ZsGreen-expressing IDLVs (integrase contained an inactivating D64V mutation) with different ratios of packaging plasmid psPAX2-D64V and psPAX2-D64V-NC-Com DNA and examined ZsGreen expression. IDLVs generated with a mixture of psPAX2-D64V and psPAX2-D64V-NC-Com DNA did not improve ZsGreen expression compared with IDLVs packaged with only psPAX2-D64V, possibly because the benefit of increased genomic RNA packaging was offset by the detrimental effects of modified Gag on gene expression. However, in IDLVs packaged with unmodified packaging plasmid psPAX2-D64V, the com sequence in the transfer plasmid evidently increased the percentage of ZsGreen-positive cells (**Fig 5B**) and their fluorescent intensity (**Fig 5C**), although particle concentrations were comparable (listed in parentheses in **Fig 5B**).

To test whether this observation reflected a vector- and gene-specific effect, we inserted an aptamer com sequence in a human *CLCN5*-expressing lentiviral transfer plasmid and prepared integration-competent LVs using packaging plasmid psPAX2. After transducing the LVs into HEK293T cells, we performed RT-qPCR using transgene-specific primers for codon optimized *CLCN5* cDNA to detect transgene expression. We found higher gene expression from aptamer-positive LVs than from aptamer-negative LVs (**Fig 5D**). Similar results occurred when the com sequence was inserted into a lentiviral transfer plasmid pFCK-HBB (n)-g1 expressing *HBB* sickle sgRNA (**Fig 5E**). These data showed that simply adding an aptamer sequence in the transfer plasmid increased gene expression of LVs.

We inserted the aptamer com outside of the ZsGreen-, *CLCN5-* or sgRNA-expression cassette; the aptamer would not be transcribed with the target gene. There was no sequence difference in the target genes whether or not the LVs had com aptamer. Thus, different target gene expression from LVs with or without com was unlikely to be caused by differences in target RNA stability.

To examine whether enhanced gene expression was the result of enhanced packaging of the LV genomic RNA into LV particles, we purified RNA from LV particles with and without the aptamer com, and performed RT-qPCR to detect the presence of the Psi sequence. Normalized with particle concentrations determined by p24 ELISA, we observed a 1- to 3-fold increase of genomic RNA in all LVs with com aptamer (**Table 2**). We and others previously reported that aptamer addition decreased the stability of host mRNA [24, 28, 48]. Thus the observed average copy number increase in LV particles could not be explained by the possible effects of com on LV genomic RNA stability. The most likely explanation was that adding the com aptamer to LV genomic RNA increased genomic RNA packaging even in the absence of ABP. This explanation was also consistent with our observation that unmodified Gag protein could also package Cas9-aptamer RNA (**Figs 3B** and **3C**; see Discussion for possible mechanisms).

**Table 2.** Effects of aptamer com on LV genomic RNA and capsid assembly.

|  | Genomic RNA copy number<sup>a</sup> |  | Viral particle concentration (ng/ml p24) |  |
| --- | --- | --- | --- | --- |
|  | No com | With com | No com | With com |
| ZsGreen LV | 2 (n=3) | 8.0 ± 0.2 (n=3) | 153.9 ± 3.6 (n=3) | 223.9 ± 1.9*** (n=3) |
| CLCN5 LV | 2 (n=3) | 6.0 ± 0.58 (n=3) | 168.2 ± 4.9 (n=6) | 293.4 ± 38.7* (n=6) |
| Sickle sgRNA LV | 2 (n=3) | 4.4 ± 1.0 (n=3) | 148.7 ± 7.9 (n=6) | 224.0 ± 20.1** (n=6) |

<sup>a</sup>Copy numbers were based on the assumption of 2 copies of genomic RNA for respective LVs without com. \*, \*\* and \*\*\* indicate p<0.05, p<0.01, and p<0.0001 respectively in two-tailed t-tests.

Assaying p24 concentrations by ELISA, we observed that including com in LV genomic RNA increased production of LVs, regardless whether they were packaged with the second generation packaging system (ZsGreen LV) or third generation packaging system (CLCN5 LV and Sickle sgRNA LV) (**Table 2**). Altogether, our data showed that including the aptamer com in LV genomic RNA increased genomic RNA packaging and capsid assembly. The increased gene expression from the modified LVs could be the results of both effects.

## 4. Discussion

Here we successfully used aptamer/ABP interactions to develop VLPs for co-delivering Cas9 mRNA and sgRNA, and added an aptamer into LV genomic RNA for improved genomic RNA packaging and particle assembly.

Delivering Cas9 mRNA or RNPs is safer than delivering Cas9 by LVs or adeno-associated viral vectors which mediate long-term Cas9 expression [18, 19]. We previously developed VLPs for Cas9 mRNA delivery [28], but they could not co-deliver sgRNA. Most other VLP-based Cas9 mRNA delivery methods also could not co-deliver sgRNA [30-32]. Fusing ABP to the N-terminus of Gag enables VLPs to co-deliver Cas9 mRNA and sgRNA expression cassettes [33, 34]. However, this method of modifying Gag decreases lentiviral particle assembly [20]. Knopp et al added RNA aptamers to both Cas9 mRNA and sgRNA and obtained inconsistent co-delivery of Cas9 mRNA and sgRNA [31]. In addition, it is possible that Cas9 RNPs rather than Cas9 mRNA plus sgRNA were delivered, as reported by Lyu et al [24] and Hamilton et al [27].

Compared with the VLP-based Cas9 mRNA and sgRNA co-delivery method reported recently [33, 34], our method reported here modified Gag with near-normal capsid assembly efficiency [24, 25, 28]. The resulting VLPs co-packaged with Cas9 mRNA and sgRNA-expressing LV genomic RNA could be used without further concentration to achieve efficient genome editing. Highly efficient particle assembly reduces costs of particle production and improves genome editing efficiency. In addition, com/Com was more efficient than the currently used MS2/MCP method for Cas9 mRNA delivery by VLPs. This observation could improve methods for delivering Cas9 for genome editing, and inform efforts to use VLPs to deliver other RNAs for various purposes.

Although VLPs can be developed to deliver Cas9 RNPs [20-27], delivering Cas9 mRNA may achieve short-term yet strong Cas9 expression to increase genome editing efficiency. This is especially relevant when targeting loci in heterochromatin regions. The nucleosome and heterochromatin inhibit Cas9 function [49-51]. This inhibitory effect is especially evident in cases of transient Cas9 expression and low Cas9 dosage [35]. Our observation that Cas9 mRNA and sgRNA all-in-one VLPs are more active in genome editing than RNP VLPs also shows the value of Cas9 mRNA and sgRNA co-delivery for efficient genome editing.

Simply including an aptamer sequence in the vector genomic RNA improved genomic RNA packaging into lentiviral capsids and particle production, which was unexpected. Currently the mechanisms by which the aptamer enhanced genomic RNA packaging is unclear. It could be caused by weak interaction between the com aptamer and unknown lentiviral proteins. Interactions between the HIV NC protein and LV genomic RNA are required for Gag assembly [52, 53]. In our experimental conditions, com increased genomic RNA packaging by 1-to 3-fold. For clinical applications, it is possible to package suitable copies of genomic RNA per particle to meet the recommended <5 copies per cell by adjusting the DNA amount of different plasmids during vector production.

In summary, this work successfully used aptamer-ABP interactions to develop VLPs for co-delivery of Cas9 mRNA and sgRNA. In addition, we showed that simply adding a com aptamer in the LV genomic RNA improved LV genomic RNA packaging and lentiviral particle assembly.

## DATA AVAILABILITY

Plasmids and sequence information are available upon request.

## SUPPLEMENTARY INFORMATION

**S1 Table**. Plasmids made for the study.

**S2 Table**. Primer sequence information for the study.

**S3 Table**. Target sequences and oligos for cloning guide into sgRNA-expressing vectors.

**S1 Fig**. Inserting Com after the second zinc finger domain greatly affected lentiviral vector titer.

**S2 Fig**. Rescuing lentiviral vector titer by unmodified package plasmid DNA.

**S3 Fig**. The GFP-expressing cassette in GFP-reporter cells.

**S4 Fig**. Efficiency of different aptamer/ABP pairs for SpCas9 mRNA delivery.

**S5 Fig**. Determining the best ratio of unmodified and modified packaging plasmid for most efficient co-packaging.

Dear Editor,

We are submitting our manuscript tilted “Developing all-in-one virus-like particles for Cas9 mRNA/single guide RNA co-delivery and aptamer-containing lentiviral vectors for improved gene expression” and authored by Manish Yadav, Anthony Atala, and Baisong Lu to your journal for evaluation. Our work develops all-in-one virus-like particles for co-delivery of Cas9 mRNA and sgRNA RNA for genome editing applications. In addition, we included aptamer in lentiviral genomic RNA to improve lentiviral vector gene expression. The data have not been previously published and are not being considered for publication elsewhere.

We hope that the manuscript be evaluated as suitable for this journal.

Sincerely,

Baisong Lu,

Associate professor

Wake Forest Medical School

