## Supplemental Tables and Figures for "Developing all-in-one virus-like particles for Cas9 mRNA/single guide RNA co-delivery and aptamer-containing lentiviral vectors for improved gene expression": SUPPLEMENTARY INFORMATION.docx

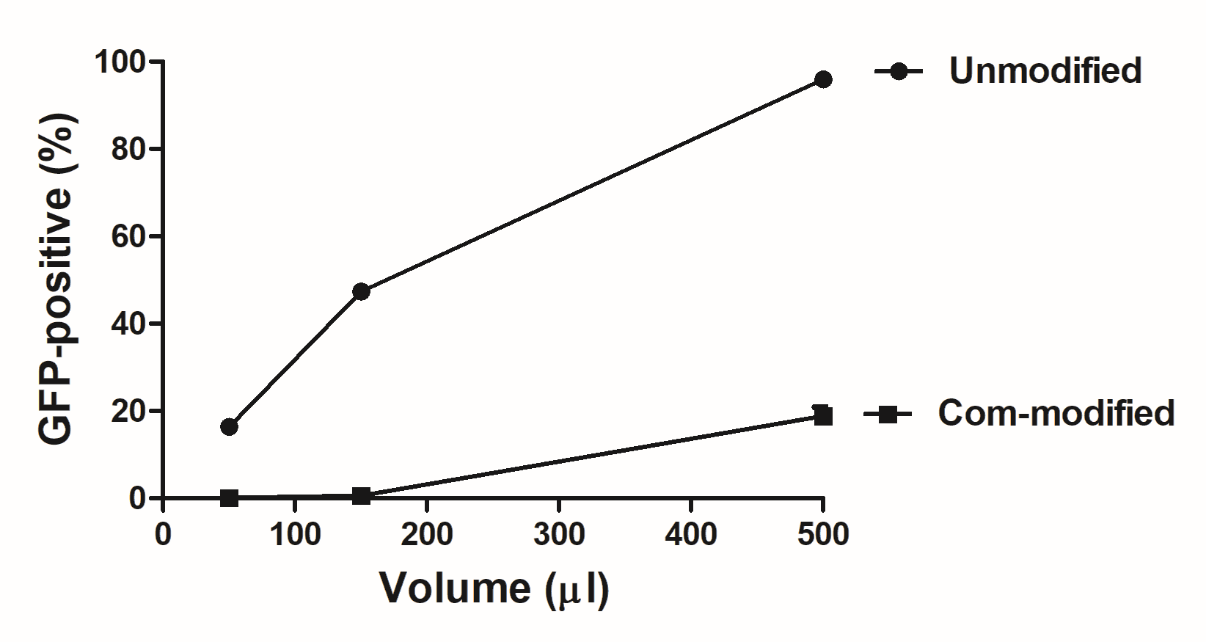

**S1 Fig.** Inserting Com after the second zinc finger domain greatly affected LV titer. The Com-modified packaging plasmid was used to generate GFP-expressing IDLVs. Each point indicates mean ± SEM of 3 technical replicates. Unmodified and Com-modified package plasmids were psPAX2-D64V and psPAX2-D64V-NC-Com, respectively.

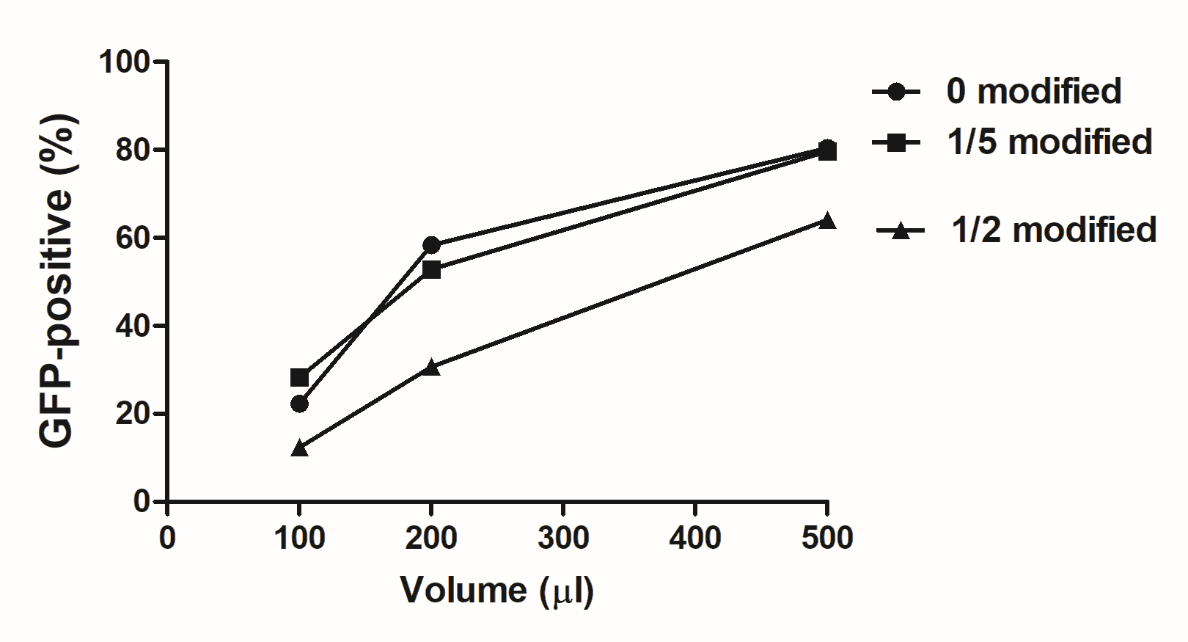

**S2 Fig.** Rescuing LV titer by unmodified package plasmid DNA. GFP-expressing IDLVs were made with different ratios of modified (psPAX2-D64V-NC-Com) and unmodified packaging plasmid DNA (psPAX2-D64V). Each point indicates mean ± SEM of 3 technical replicates. Unmodified and Com-modified package plasmids were psPAX2-D64V and psPAX2-D64V-NC-Com, respectively.

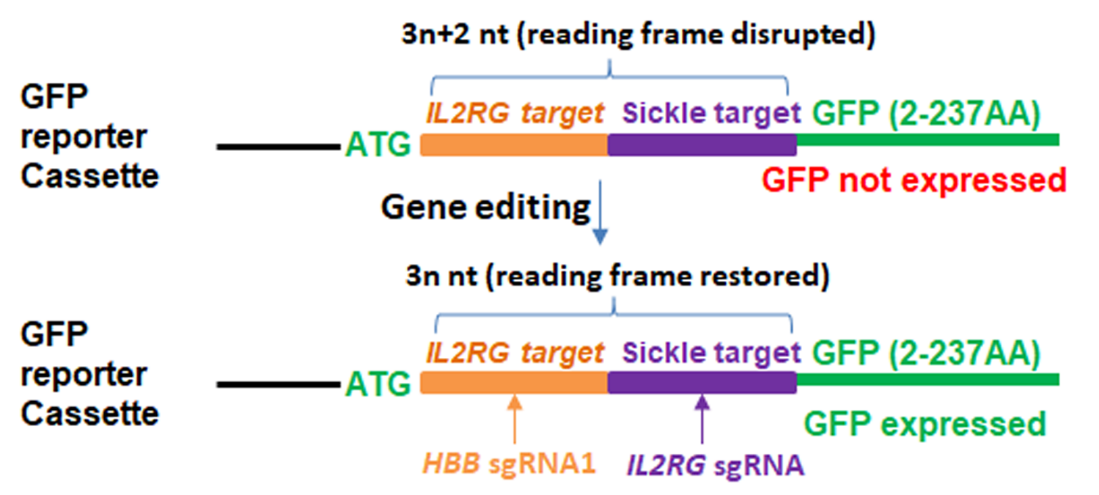

**S3 Fig.** The GFP expression cassette in GFP-reporter cells. Insertions and deletions in target sites may restore the GFP reading frame and GFP expression.

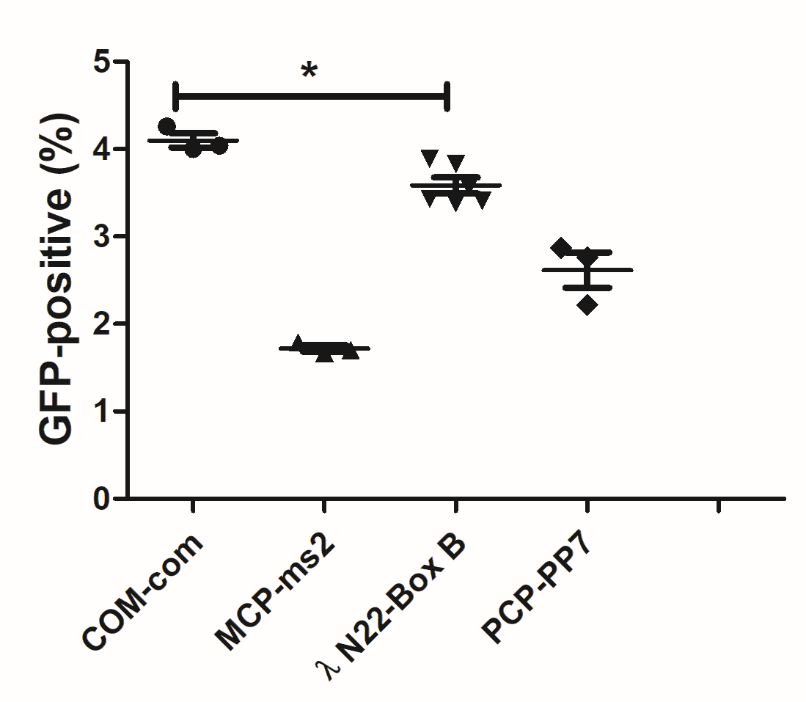

**S4 Fig.** Efficiency of different aptamer/ABP pairs for SpCas9 mRNA delivery. Plasmid DNA expressing *IL2RG* sgRNA was transfected into the GFP-reporter cells 12 hours before transduction of 500 µl SpCas9 mRNA-packaged VLPs. * indicates p<0.05 for com/Com VLPs versus BoxB/λN22 VLPs (Tukey’s *post hoc* analysis following ANOVA).

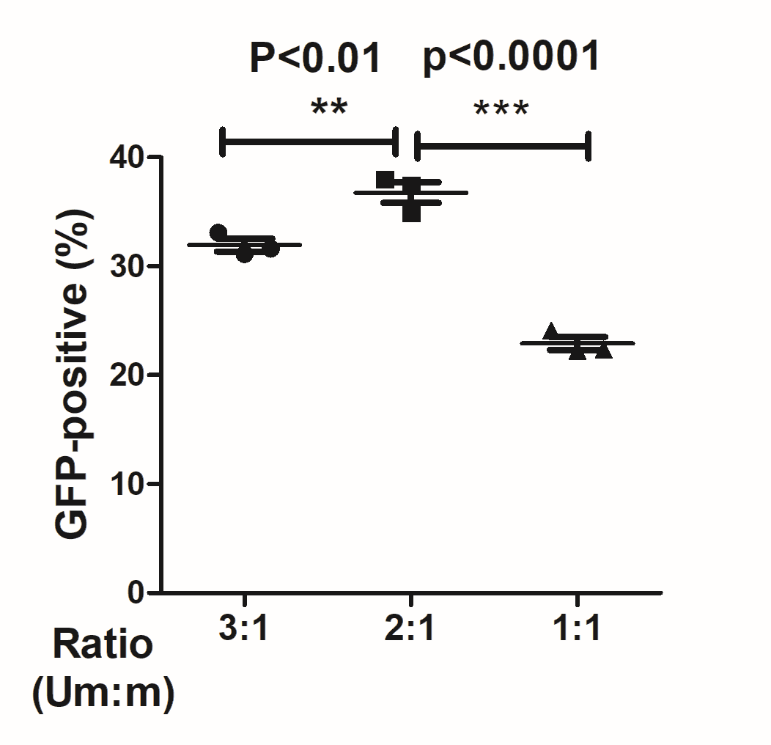

**S5 Fig.** Determining the best ratio of unmodified and modified packaging plasmid for most efficient co-packaging. Indicated ratios of unmodified (Um) packaging plasmid (psPAX2-D64V) and Com-modified (M) packaging plasmid (psPAX2-D64V-NC-Com) were used to transfect HEK293T cells by FuGENE HD to make VLPs, which contained Cas9 mRNA and LV genomic RNA for expressing sickle sgRNA. The particles (400 µl) were transduced into GFP-reporter cells and GFP-positive cells were analyzed by flow cytometry. Each point indicates mean ± SEM of 3 biological replicates.

**S1 Table . Plasmids made for the study**

| No. | Name | Purpose | Generation strategy |
| --- | --- | --- | --- |
| 1 | pBlue-SaCas9-3 aptamers | A plasmid-expressing SaCas9 protein with three aptamers (com, PP7, and boxB) and two copies of the human beta globin gene 3’ UTR after the stop codon of SaCas9. | A synthesized DNA fragment “ttttcattgcgaattcACTGAATGCCTGCGAGCATCCCACtcgcgaggagcagacgatatggcgtcgctccGTTAACGGGCCCTGAAGAAGGGCCCtcaggtcgactctagaaaat” containing com, PP7, and boxB aptamer sequences was inserted into the EcoRI and SalI sites of pX601-ms2-3’UTR by infusion cloning. The BamHI-EagI fragment from pX601-ms2-3’UTR was inserted into the BamHI-EagI sites of pBlue-Cas9-tetra-com-sickle-g1 to obtain pBlue-Sacas9-3 aptamers. |
| 2 | pBlue-spCas9-3 aptamers | A plasmid-expressing SpCas9 protein with three aptamers (com, PP7, and boxB) and two copies of human beta globin gene 3’ UTR after the stop codon of SpCas9. | The SaCas9 sequence in pBlue-SaCas9-aptamers was replaced with SpCas9 sequences by restriction and ligation. |
| 2 | psPAX2-D64V-COM-NC | Packaging plasmid with COM fused to the N-terminus of NC | DNA fragment 1 was amplified with Sph1-F and COM-NC-MR1 from psPAX2-D64V, DNA fragment 2 was amplified with COM-NC-MF1 and COM-NC-MR2 from psPAX2-D64V-NC-COM, and DNA fragment 3 was amplified with COM-NC-MF2 and COM-NC-R from psPAX2-D64V. The three fragments were used as templates for overlapping PCR. PCR products were inserted into the SphI and SbfI sites by infusion cloning. |
| 3 | psPAX2-D64V-COM | Packaging plasmid with COM inserted between NC and P6. Protease cleavage sites were included to produce intact NC and P6 proteins after processing. | DNA fragment 1 was amplified with Sph1-F and COM-inMR from psPAX2-D64V-NC-com, DNA fragment 2 was amplified with COM-inMF and COM-inR from psPAX2-D64V-NC-com. The two fragments were used as templates for overlapping PCR. PCR products were inserted into the SphI and BmgBI sites by infusion cloning |
| 4 | pLH-sgRNA-IL2RG | Lentiviral transfer plasmid expressing IL2RG targeting sgRNA for SpCas9. | Annealed oligos of Il2RG-sp-g1R ( AAACAGGGAATGAAGAGCAAGCGC) and Il2RG-sp-g1F1 (ACCGGCGCTTGCTCTTCATTCCCT) were inserted into the Bbs1 site of pLH-sgRNA1 by T4 DNA ligase. |
| 5 | pFCK-HBB(n)-g1 | Lentiviral transfer plasmid expressing sickle mutant sequence targeting sgRNA for SaCas9. | See [38]. |
| 6 | pFCK-com-HBB(n)-g1 | Lentiviral transfer plasmid expressing sickle mutant sequence targeting sgRNA for SaCas9. Aptamer com was inserted in KflI site of the plasmid. | Annealed oligos from Com for LV-F1 ( caaccccgaggggacCTGAATGCCTGCGAGCATCCCACgacccgacaggcccg) and Com for LV-R1 ( cgggcctgtcgggtcGTGGGATGCTCGCAGGCATTCAGgtcccctcggggttg) were inserted into the KflI site of pFCK-com-HBB(n)-g1 by infusion reaction. |
| 7 | pCSII-hCLCN5 | Lentiviral transfer plasmid expressing human CLCN5 cDNA. | Condon optimized human CLCN5 cDNA sequence (ctcgagccaccATGGATTTCCTgGAGGAACCAATACCAGGTGTAGGAACATATGACGATTTCAATACTATAGACTGGGTGCGAGAGAAATCACGCGATCGAGACAGACACCGGGAGATCACGAATAAGTCTAAGGAATCTACCTGGGCCCTCATTCACAGTGTGTCAGACGCTTTTAGCGGATGGCTGCTTATGCTTCTGATTGGACTTCTTAGTGGTAGTTTGGCGGGCCTGATAGACATTAGCGCGCACTGGATGACTGATCTTAAAGAAGGCATATGCACGGGGGGATTTTGGTTCAACCACGAACATTGCTGCTGGAACTCCGAGCATGTGACATTCGAGGAGAGGGACAAGTGCCCCGAGTGGAATAGTTGGAGCCAACTGATAATTTCTACAGATGAGGGGGCTTTTGCCTATATAGTTAATTATTTCATGTATGTTTTGTGGGCCCTCCTCTTCGCCTTCCTCGCGGTATCCCTCGTTAAGGTCTTTGCCCCATATGCCTGTGGCTCTGGTATTCCAGAAATAAAAACTATCCTTTCTGGATTTATAATCAGGGGATATCTGGGCAAGTGGACGTTGGTCATTAAGACAATCACCCTTGTCCTTGCTGTATCTTCAGGGTTGTCCTTGGGCAAAGAGGGTCCTCTCGTTCACGTAGCTTGCTGCTGTGGGAACATCCTTTGCCATTGTTTCAATAAATATAGGAAGAACGAAGCAAAGCGCCGAGAAGTTCTGAGCGCAGCAGCGGCCGCAGGTGTCAGTGTTGCCTTCGGGGCTCCTATAGGAGGGGTACTGTTTAGTCTCGAAGAAGTGTCATATTACTTTCCTCTCAAGACACTGTGGAGGTCCTTTTTTGCAGCCCTGGTCGCGGCTTTTACTCTGCGCTCTATTAATCCTTTTGGAAACAGCAGACTTGTGCTGTTCTACGTCGAATTCCACACCCCGTGGCATTTGTTTGAACTCGTACCCTTTATTTTGCTGGGGATTTTCGGTGGATTGTGGGGTGCTCTGTTCATACGCACTAACATTGCGTGGTGCCGGAAGAGGAAGACTACTCAGTTGGGCAAATACCCAGTTATTGAGGTCCTCGTCGTTACAGCTATCACAGCAATTCTTGCGTTCCCCAACGAGTACACACGGATGTCTACATCCGAACTGATTAGCGAACTGTTCAATGATTGTGGGCTCTTGGACTCCTCAAAACTGTGCGATTATGAAAATCGATTTAATACATCAAAGGGCGGAGAACTTCCCGATCGGCCGGCTGGAGTGGGAGTATACTCCGCTATGTGGCAGCTGGCGTTGACGCTCATACTCAAAATCGTCATTACCATATTCACTTTTGGAATGAAGATTCCCTCAGGTCTCTTTATCCCTAGTATGGCAGTTGGTGCGATTGCGGGACGGCTCCTGGGCGTTGGCATGGAGCAGCTGGCTTATTACCATCAGGAGTGGACCGTATTCAATAGCTGGTGCTCTCAGGGCGCTGATTGCATCACACCAGGCCTGTATGCCATGGTAGGCGCTGCTGCTTGTCTTGGAGGGGTGACTAGGATGACGGTTTCTCTCGTCGTGATAATGTTCGAGCTTACTGGGGGTCTTGAGTACATTGTGCCCCTGATGGCGGCGGCAATGACATCCAAATGGGTGGCGGATGCGTTGGGTAGGGAAGGGATATACGATGCACATATTCGCCTTAATGGCTACCCATTTTTGGAGGCTAAGGAAGAATTTGCACATAAAACTCTCGCCATGGATGTTATGAAACCGAGACGAAACGACCCATTGCTTACAGTACTTACACAGGATTCCATGACCGTTGAGGACGTGGAAACAATAATATCTGAAACAACTTATAGTGGCTTTCCCGTCGTCGTATCCCGAGAATCACAAAGGTTGGTAGGATTCGTGCTGCGACGCGACCTGATCATATCCATAGAAAACGCACGCAAGAAGCAAGACGGGGTAGTGTCCACGTCTATAATTTATTTCACCGAGCATAGCCCTCCCTTGCCTCCATATACTCCGCCTACACTGAAACTTCGAAACATCCTCGATTTGTCTCCTTTTACAGTAACCGACCTTACTCCAATGGAAATCGTAGTAGACATATTTAGAAAGCTTGGATTGAGGCAATGCCTGGTTACCCACAACGGTCGGTTGCTCGGGATAATAACGAAGAAGGACGTACTCAAACATATAGCACAAATGGCAAACCAGGACCCgGATTCAATCTTGTTCAACTAGtctaga) was synthesized by GenScript. The DNA fragment was inserted between XhoI and XbaI sites of pCSII-EF-miRFP709-hCdt (from Addgene). |
| 8 | pCSII-com-hCLCN5 | Lentiviral transfer plasmid expressing human CLCN5 cDNA. Aptamer com was inserted in KflI site of the plasmid. | The annealed oligos from Com for LV-F1 ( caaccccgaggggacCTGAATGCCTGCGAGCATCCCACgacccgacaggcccg) and Com for LV-R1 ( cgggcctgtcgggtcGTGGGATGCTCGCAGGCATTCAGgtcccctcggggttg) were inserted into the KflI site of pCSII-hCLCN5 by Infusion reaction. |
| 9 | pLVX-IRES-ZsGreen1 | Lentiviral transfer plasmid expressing ZsGreen. | Ordered from Takara Inc. |
| 10 | pLVX-com-IRES-ZsGreen1 | Lentiviral transfer plasmid expressing ZsGreen. Aptamer com was inserted in KflI site of the plasmid. | The annealed oligos from Com for LV-F1 ( caaccccgaggggacCTGAATGCCTGCGAGCATCCCACgacccgacaggcccg) and Com for LV-R1 ( cgggcctgtcgggtcGTGGGATGCTCGCAGGCATTCAGgtcccctcggggttg) were inserted into the KflI site of pLVX-IRES-ZsGreen1 by Infusion reaction. |
| 11 | pSaCas9^1xcom^-2x3’UTR | Two copies of 3’ untranslated region (UTR) from human *HBB* gene were inserted after the Sacas9 coding sequences and before the com aptamer. | The MS2 aptamer of pSaCas9^1xMS2^-2x3’UTR was replaced with the com aptamer with a sequence of ACTGAATGCCTGCGAGCATCCCAC. |
| 12 | pSpCas9^1xMS2^-2x3’UTR | Two copies of 3’ untranslated region (UTR) from human *HBB* gene were inserted after the Spcas9 coding sequences and before the MS2 aptamer. | The annealled products of oligonucleotides sp-loop1F (AAAAAAGAAAAAGCTTTAGAAAACATGAGGATCACCCATGTCTGCAGGTCGACTCTAGAATTCCTAGAGCTCG) and sp-loop1R (CGAGCTCTAGGAATTCTAGAGTCGACCTGCAGACATGGGTGATCCTCATGTTTTCTAAAGCTTTTTCTTTTTT) was inserted into the HindIII and EcoR1 sites of pU6-sgRosa26-1_CBh-Cas9-T2A-BFP (Addgene #64216) to obtain pSpCas9^1xMS2^. Then the FseI-EagI fragment containig two copies of the HBB 3’ UTR sequences were inserted into the FseI and NotI sites of pSpCas9^1xMS2^. |
| 13 | pSpCas9^1xcom^-2x3’UTR | Two copies of 3’ untranslated region (UTR) from human *HBB* gene were inserted after the Spcas9 coding sequences and before the com aptamer. | The BamHI-PvuI fragment of pSaCas9^1xcom^-2x3’UTR was inserted between the BamHI-PvuI sites of pSpCas9^1xMS2^-2x3’UTR to replace the MS2 aptamer with the com aptamer. |

**S2 Table. Primers used for the study**

| Primer name | SEQ | Use |
| --- | --- | --- |
| hCLCN5-CF | GAAGCAAGACGGGGTAGTGTC | To detect CLCN5 expression from transgene by qPCR |
| hCLCN5-CR | GGTCGGTTACTGTAAAAGGAGACAA |  |
| Sickle-F1 | CACCGAGTAACGGCAGACTTCTCCAC | To detect sickle sgRNA expression by qPCR |
| sgRNA-R3 | GATAAACACGGCATTTTGCCTTG |  |
| Psi-F | TCTCGACGCAGGACTCG | To detect LV genomic RNA |
| Psi-R | TACTGACGCTCTCGCACC |  |
| Sph1-F | agagtgcatccagtgcatgcag | To amplify DNA for making psPAX2-D64V-COM-NC |
| COM-NC-MR1 | atggatccacctccaccggaattgcctttctgtatcattatgg | To amplify DNA for making psPAX2-D64V-COM-NC |
| COM-NC-MF1 | ccataatgatacagaaaggcaattccggtggaggtggatccat | To amplify DNA for making psPAX2-D64V-COM-NC |
| COM-NC-MR2 | cctaaaattgcctttctgtattccacctccaccTCCGGAGTTGTGACCGCCataacgcacggtttc | To amplify DNA for making psPAX2-D64V-COM-NC |
| COM-NC-MF2 | atacagaaaggcaattttaggaacc | To amplify DNA for making psPAX2-D64V-COM-NC |
| COM-NC-R | tttctgttttaaccctgcaggatg | To amplify DNA for making psPAX2-D64V-COM-NC |
| COM-inMR | aAatTttAccAAGGaaGttTgcTtgACGTtcagtacaatctttcatttggtgtc | To amplify DNA for making psPAX2-D64V-COM |
| Com-inMF | GTcaAgcAaaCttCCTTggTaaAatTtccggtggaggtggatccatggct | To amplify DNA for making psPAX2-D64V-COM |
| COM-inR | AATTGATTTCATCACgtcgccagt | To amplify DNA for making psPAX2-D64V-COM |

**S3 Table. Target sequences and oligonucleotides for cloning the guide sequences into sgRNA-expressing vector**

| Target gene | Target sequence with PAM | Forward Oligo for cloning | Reverse Oligo for cloning | Plasmid name |
| --- | --- | --- | --- | --- |
| *IL2RG* | GCGCTTGCTCTTCATTCCCTGGG | ACCGGCGCTTGCTCTTCATTCCCT | AAACAGGGAATGAAGAGCAAGCGC | pLH-sgRNA1-2xMS2-IL2RG |
| *HBB* Sickle mutant sequence | AGTAACGGCAGACTTCTCCTCAGGAGT | CACCGAGTAACGGCAGACTTCTCCAC | AAACGTGGAGAAGTCTGCCGTTACTC | pFCK-HBB(n)-g1 |
